# A high-resolution diel survey of surface ocean metagenomes, metatranscriptomes, and transfer RNA transcripts

**DOI:** 10.1101/2025.09.15.676277

**Authors:** Sarah J. Tucker, Jessika Fuessel, Kelle C. Freel, Evan Kiefl, Evan B. Freel, Oscar Ramfelt, Clarisse E. S. Sullivan, Andrian P. Gajigan, Hanako Mochimaru, Mariana Rocha de Souza, Mariko Quinn, Ciara Ratum, Leon L. Tran, Marek Sobczyk, Samuel E. Miller, Florian Trigodet, Karen Lolans, Hilary G. Morrison, Bailey Fallon, Bruno Huettel, Tao Pan, Michael S. Rappé, A. Murat Eren

## Abstract

The roles of marine microbes in ecosystem processes are inherently linked to their ability to sense, respond, and ultimately adapt to environmental change. Capturing the nuances of this perpetual dialogue and its long-term implications requires insight into the subtle drivers of microbial responses to environmental change that are most accessible at the shortest scales of time. Here, we present a multi-omics dataset comprising surface ocean metagenomes, metatranscriptomes, tRNA transcripts, and biogeochemical measurements, collected every 1.5 hours for 48 hours at two stations within coastal and adjacent offshore waters of the tropical Pacific Ocean. We expect that this integrated dataset of multiple sequence types and environmental parameters will facilitate novel insights into microbial ecology, microbial physiology, and ocean biogeochemistry and help investigate the different mechanisms of adaptation that drive microbial responses to environmental change.

## Background & Summary

Microbial communities experience environmental fluctuations over a broad range of time-scales, and their responses are evident at multiple levels – from changes in community composition to the physiological reactions of individual cells^1^. Time-series studies of marine microbes have largely focused on time-scales of weeks, months, and seasons^2–8^, with a particular emphasis on changes to microbial community composition. While these studies have provided critical insights into marine microbial biogeography, diversity, and ecology, the physiological shifts of individual microbial populations, especially at short timescales, are less well understood.

Microbes can rapidly respond to environmental change via transcriptional and translational regulation, often without any immediate impact on community composition^9,10^. These changes enable rapid functional adaptations that modulate microbial metabolisms^11^, influence microbial contributions to ocean biogeochemical cycles^12^, shape biogeographic distributions^13^, and affect long-term ecological and evolutionary trajectories^14^. Yet, relatively few studies have sampled gene expression and community composition with high enough temporal resolution to track the reactions of microbial populations to environmental fluctuations and the activities of other community members^9,15–17^.

Here, we present a multi-omics dataset from our recent Hawaiʻi Diel Sampling (HaDS) survey, which represents a high-resolution sampling of surface ocean microbial communities every 1.5 hours for a two-day period across two environments: Kāneʻohe Bay on the windward coast of Oʻahu, Hawaiʻi, and the adjacent offshore (**Fig. 1a,b**). The goal of HaDS is to characterize microbial responses to diel changes in ocean biogeochemistry across physically connected, yet environmentally distinct habitats. We designed HaDS to focus on sampling stations previously established within the Kāneʻohe Bay Time-series (KByT)^18,19^, so that the diel data can be further contextualized with insights from monthly observations

**Figure 1.**
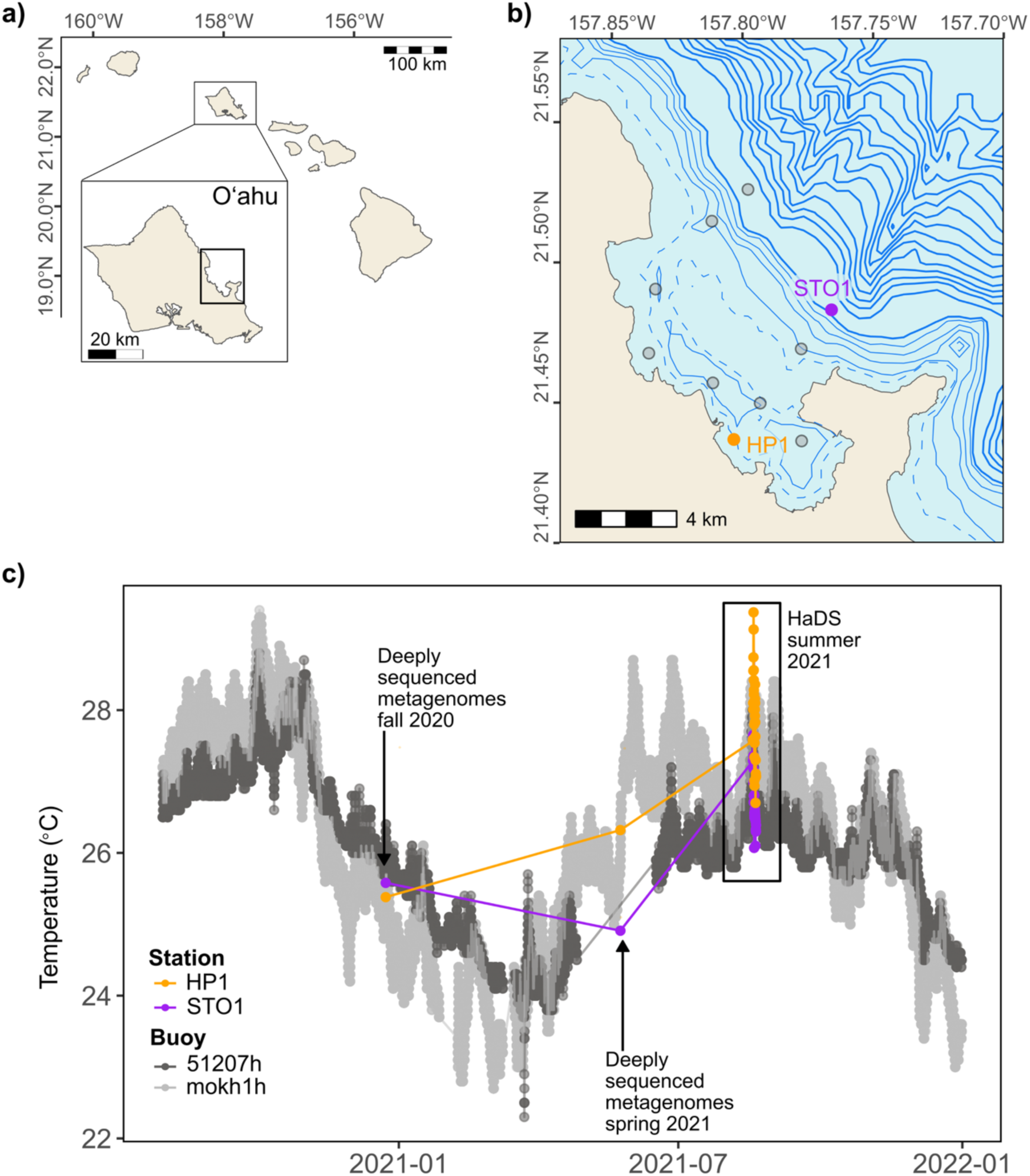
Sampling regime of HaDS. **a)** Map showing the location of Kāneʻohe Bay on the windward side of the island of Oʻahu, Hawaiʻi. **b)** We sampled stations HP1 (coastal Kāneʻohe Bay) and STO1 (adjacent offshore) during HaDS. These stations are part of the monthly sampling of the Kāneʻohe Bay Time-series (KByT), which also includes 8 additional KByT sampling stations shown in gray. Dashed bathymetric contours mark a depth of 5 m. Solid bathymetric contours indicate depth increases of 10 m up to 50 m, with thicker lines marking 50 m depth intervals. **c)** We conducted HaDS sampling during a period of elevated summer temperatures. Two additional sampling events contributed four deeply sequenced metagenomes that contextualize the summer diel sampling. Temperature data collected during the sampling events and from buoys located in coastal Kāneʻohe Bay (mokh1h; National Buoy Data Center) and the adjacent offshore (51207h; National Buoy Data Center) show differences in surface water temperature between coastal and offshore stations as well as seasonal variation throughout the data collection period.

Altogether, HaDS generated 202 ‘omics sequencing products totalling 6.62 billion paired-end short reads and 27.63 million single-end long reads that capture multi-dimensional, high-resolution observations of nearshore and offshore microbial communities (**Table S1**). At both the coastal Kāneʻohe Bay station (HP1) and the adjacent offshore station (STO1), we sampled at 33 time points across 48 hours, and subsequently produced 59 metatranscriptomes, 65 short-read metagenomes, 8 long-read metagenomes, and 66 transfer RNA (tRNA) transcript libraries (**Table 1**), as well as biogeochemical measures. To the best of our knowledge, HaDS includes the first high-throughput sequencing of transfer RNA transcripts from marine microbial communities, offering a unique dataset for analysis. We also generated four deeply-sequenced short-read metagenomes from samples collected in the late fall and spring prior to HaDS through routine KByT sampling (**Fig. 1c**, **Table 1**).

**Table 1.** Summary of multi-omics datasets collected as part of the Hawaiʻi Diel Sampling (HaDS) and the Kāneʻohe Bay Time-series (KByT).

| Project | Station | Sampling dates | Short-read metagenomes | Long-read metagenomes | Metatranscriptomes | tRNA transcripts |
| --- | --- | --- | --- | --- | --- | --- |
| HaDS | HP1 | 8/18/21-8/20/21 | 32 | 4 | 29 | 33 |
| HaDS | STO1 | 8/18/21-8/20/21 | 33 | 4 | 30 | 33 |
| KByT | HP1 | 5/24/21 | 1 | - | - | - |
| KByT | STO1 | 5/24/21 | 1 | - | - | - |
| KByT | HP1 | 12/23/20 | 1 | - | - | - |
| KByT | STO1 | 12/23/20 | 1 | - | - | - |

In addition to the high-quality, raw sequencing data and contextual measures of the environment, we provide station-specific co-assemblies based on both the short-read and long-read metagenomes that were collected during the diel sampling. We reconstructed a total of 160 metagenome-assembled genomes (MAGs) from the short-read metagenomes, including 63 high-quality (>90% completeness, <5% contamination) and 97 medium-quality (>70% completeness, <10% contamination) MAGs (**Table S1**). Co-assemblies of long-read metagenomes per-station yielded 73 contigs over 1 Mbp, 14 high-quality MAGs, and 16 medium-quality MAGs. By providing the reconstructed MAGs, co-assemblies for short- and long-read metagenomes, and associated workflows in addition to the raw data, we aim to reduce the computational burden associated with this analysis step and ultimately increase the accessibility of these datasets for the broader research community. However, future analyses should consider multiple assembly approaches including hybrid assembly^20^ and eukaryotic-specific assemblers^21^, as we expect that these analyses may capture additional populations potentially missed in our co-assemblies.

In summary, the Hawaiʻi Diel Sampling effort provides novel and cutting-edge multi-omics data alongside complementary environmental parameters, metagenomic assembly data products, and reproducible bioinformatic workflows that enhance the utility and value of these unique resources.

## Methods

### Hawaiʻi Diel Sampling (HaDS) collection and processing

Seawater collection took place between August 18, 2021 9:00 AM and August 20, 2021 9:00 AM Hawaiʻi Standard Time (HST), from a depth of ∼0.25 m at station HP1 within coastal Kāneʻohe Bay, Oʻahu, Hawaiʻi, and station STO1 in the adjacent offshore waters. Our periodic sampling every 1.5 hours over a 48 hour period yielded 33 sampling events at each site. For each time point, we collected seawater samples for flow cytometry, fluorometric chlorophyll *a* concentrations, and nucleic acids. Additionally, we recorded *in situ* measurements of seawater temperature, pH, and salinity with a YSI 6600 or YSI EcoSense EC300 (YSI Incorporated, Yellow Springs, OH, USA) and *in situ* measurements of Photosynthetic Photon Flux Fluence Rate (PPFFR) with a LI-193 Underwater Spherical Quantum Sensor and LI-250A Light Meter (LI-COR Environmental, Lincoln, NE, USA).

Following prefiltration of ∼10-20 L of seawater through 85-μm Nitex mesh, we filtered ∼4 (HP1) or ∼9 (STO1) L of seawater through 0.2-μm pore-sized polyethersulfone (PES) Sterivex cartridge filters (MilliporeSigma, Burlington, MA, USA) to collect microbial cells for nucleic acids isolation. We immediately added RNAlater (Invitrogen, Waltham, MA, United States) to the Sterivex filters and stored the samples at −80 °C until further processing.

We collected seawater subsamples (125 mL) on 25-mm diameter GF/F glass microfiber filters (Whatman, GE Healthcare Life Sciences, Chicago, IL, USA) and stored the filters in aluminum foil at −80 °C until fluorometric analysis of chlorophyll *a* concentrations. We extracted chlorophyll *a* in 100% acetone and determined the concentration with a Turner 10-AU fluorometer (Turner Designs, Sunnyvale, CA, USA) according to established techniques^22^. We preserved seawater for cellular enumeration in 2 mL aliquots with a final concentration of 0.95% (v:v) paraformaldehyde (Electron Microscopy Services, Hatfield, PA, USA) and stored the samples at −80 °C until analysis. The SOEST Flow Cytometry Facility performed flow cytometric cellular enumeration of non-cyanobacterial (presumably heterotrophic) bacteria and archaea (reported here as heterotrophic bacteria), photosynthetic picoeukaryotes, *Synechococcus*, and *Prochlorococcus* on a Beckman Coulter CytoFLEX S, following the method of Monger & Landry^23^. The SOEST Laboratory for Analytical Biogeochemistry (S-LAB) measured concentrations of dissolved inorganic nutrients on a Seal Analytical AA3 HR Nutrient Autoanalyzer (detection limits: NO_2−_ + NO_3−_, 0.009 µM; SiO_4_ 0.09 µM; PO_4_^3−^ 0.009 µM; NH_4_ 0.03 µM) and dissolved organic carbon and total nitrogen on a Shimadzu High-Temperature TOC-L Combustion Analyzer.

We used various monitoring platforms to collect additional environmental data. We accessed surface ocean temperature data covering the period from 2020–2021 from the NOAA station mokh1 in Kāneʻohe Bay and surface ocean temperatures and wave height from the NOAA station 51207 in the adjacent offshore waters. We accessed rainfall and wind speed from the NOAA Moku o Loʻe Weather station and tidal height from the NOAA/NOS/CO-OPS station 1612480, both of which are positioned on the island of Moku o Loʻe in southern Kāneʻohe Bay, along with the Hawaiʻi Institute of Marine Biology. We report tidal height as the mean lower low water, or the average of the lower low water height of each tidal day examined over the National Tidal Datum Epoch. We downloaded these data from their respective websites between November 20, 2024 and February 26, 2025.

### DNA extraction, preparation, and sequencing for short-read metagenomics

We adapted a previously published protocol^24^ to extract DNA from Sterivex cartridge filters for short-read metagenomic sequencing. First, we collected any cells suspended in the RNAlater contained in the Sterivex filter cartridges by attaching a 10 mL air-filled clean syringe to the Sterivex cartridge and pushing the liquid into a nuclease-free 1 mL centrifuge vial. After centrifugation for 15 min at 4 **°**C at 13,000 rpm, we discarded the supernatant and kept the cell pellet on ice until further processing. We then opened the cartridge holding the filter with a sterilized PVC pipe cutter and removed the cylinder with the PES membrane. Using a sterilized scalpel, we cut along the barrel at the top and bottom of the filter membrane and down one side to remove the filter from the barrel. We cut the filter into ∼6-8 small pieces in a sterile petri dish and transferred the pieces to a Lysing Matrix E vial (MP Biomedicals, Irvine, CA, USA) and added 450 µL Buffer A, 210 µL 20% SDS and 500 µL phenol:chloroform:IAA 125:24:1 pH 8 (50 mL Buffer A: 2 mL 5 M NaCl, 2 mL 0.5 M EDTA, 46 mL dH2O, all solutions were molecular grade and nuclease-free). We resuspended the cell pellet obtained from RNAlater in 50 µl Buffer A by vortexing vigorously for 30 sec followed by a quick centrifugation step in a microcentrifuge and transferred the solution to the Lysing Matrix E vial. We lyzed the cells with a Bead Ruptor Elite (OMNI International, Kennesaw, GA, USA) on setting 6 for 30 sec followed by two min on ice to avoid overheating of the samples and another lysing cycle with the same settings. We centrifuged the vials for 3 min at 4 **°**C at 13,000 rpm and removed the upper aqueous phase to a fresh, nuclease-free 1.5 mL centrifuge vial. Following the addition of 500 µL phenol:chloroform:IAA, we repeated the centrifugation step. Following transfer of the aqueous phase to a new centrifuge vial, we added 3 M sodium acetate at 1/10th of the transferred volume and mixed the solution by inversion. We added 500 µL ice-cold pure ethanol and placed the samples in the freezer at -20 **°**C overnight.

The next day, we thawed the samples on ice, centrifuged them at 13,000 rpm at 4 **°**C for 20 min, and carefully decanted the supernatant to retain the nucleic acid pellet. We washed the pellet with 500 µL 70% ethanol at 4 **°**C and vortexed shortly followed by centrifugation at 13,000 rpm at 4 **°**C for 10 min. We decanted the liquid carefully, dried the lip of the tube on a clean tissue, and air-dried the pellet with the centrifuge tubes open and upside down on a clean tissue for 15 min. We added 100 µL of nuclease-free dH2O to the pellet and vortexed the pellet for 20 min with a Vortex-Genie 2 (Scientific Industries, Bohemia, NY, USA) and a Vortex adapter 24 (Qiagen, Hilden, Germany). To remove any impurities and chemical contaminations, we purified the DNA with the DNA Clean & Concentrator-100 kit (Zymo Research, Irvine, CA, USA) and eluted in a final volume of 30 µL.

Following shearing of DNA to 400 bp on a Covaris S220 focused-ultrasonicator system, we cleaned and concentrated the DNA with AMPure XP beads (Beckman Coulter, Brea, CA, USA) and prepared metagenomic sequencing libraries using the Ovation Ultralow System V2 with indexed sequencing adapters (Tecan Genomics, Männedorf, Switzerland). We pooled final libraries at equimolar amounts and size-selected the library using a BluePippin 1.5% agarose gel cassette (Sage Science, Beverly, MA, USA) targeting the 468–572 bp region. We paired-end sequenced the final pool on a NextSeq 500/550 High Output v2.5 using the 300 cycles kit (Illumina, San Diego, CA, USA). The Keck Sequencing Facility at the Marine Biological Laboratory in Woods Hole, MA, USA carried out the short-read metagenomic library preparation and sequencing.

### Nucleic acid extraction, preparation, and sequencing for long-read metagenomics and metatranscriptomics

To extract nucleic acids from the Sterivex cartridge filters, we first removed the RNAlater by attaching a 10 mL air-filled clean syringe to the Sterivex cartridge to remove and discard the liquid. We then opened the cartridge holding the filter with a sterilized PVC pipe cutter and removed the cylinder with the PES membrane. Using a sterilized scalpel, we cut along the barrel at the top and bottom of the filter membrane and down one side to remove the filter from the barrel. We cut the filter into ∼6-8 small pieces in a sterile petri dish and transferred them to 2 mL BashingBead lysis tubes with 0.1 & 0.5 mm beads (Zymo Research Europe, Germany) for subsequent cell disruption with a Qiagen TissueLyser (3 min. at 25 Hz). Next, we isolated DNA and total RNA from the same starting material using the ZymoBIOMICS DNA/RNA Miniprep kit (Zymo Research Europe, Germany), with an additional DNase I treatment of the extracted RNA as recommended by the vendor.

DNA directly served as input for a barcoded DNA library preparation without prior fragmentation. We prepared HiFi SMRTbell libraries (PacBio, Menlo Park, CA, USA) for long-read DNA sequencing following manufacturer protocols using the Ultra Low DNA Input approach and sequenced the libraries on a PacBio Revio device. For demultiplexing, adapter trimming, and deduplication of the sequencing data we relied on the PacBio SMRTlink software suite.

We prepared RNAseq libraries with the Universal Prokaryotic RNA-Seq Prokaryotic Any Deplete kit by Tecan Genomics, including the proprietary adapter addition step of the kit and dual 8-mer barcode indexing. We sequenced the libraries on an Illumina NextSeq2000 platform in 150 bp paired-end read mode. The MPI Cologne Genomics Center conducted nucleic acid extraction, library preparation, and sequencing for long-read metagenomics and metatranscriptomics.

### RNA extraction and preparation for transfer RNA sequencing

To maximize recovery of total RNA from the Sterivex filter cartridges, we performed pre-processing steps that enhance bacterial cell lysis. For enzymatic cell lysis, we prepared the following solutions: 1) marine microbe lysis solution: 40 mM EDTA, 50 mM TRIS and 0.75 M sucrose final concentration, 2) lysozyme stock solution: 100 mg/mL in TE buffer pH 8, RNAse free, and 3) a final lysis solution of 670 µL per sample, consisting of 600 µL marine microbe lysis solution, 67 µL lysozyme stock solution, 0.2 µL SUPERase (Invitrogen™, Thermo Fisher Scientific, Waltham, MA, USA, 20 U/μL).

For each Sterivex cartridge, we collected and pelleted cells that were suspended in RNAlater following the same steps described above. We resuspended each pellet in 70 µL of lysis solution. We removed the filter from the Sterivex cartridges using the same protocol described above, cut the filter into small pieces using a clean scalpel, and transferred the pieces to a 14 mL culture tube containing 600 µL of pre-prepared lysis solution. We added the resuspended cell lysate to each 14 mL tube that contained the corresponding filter pieces and vortexed vigorously for 2 min. We then incubated the tubes in a 37 °C water bath for 45-60 min and gently agitated the samples every 10 min.

After the initial incubation, we added 7.4 µL of proteinase K (800 U/mL, New England Biolabs, Ipswich, MA, USA) and 74 µL of 10% SDS for a final concentration of 1% to each sample and incubated at 37 °C for an additional 2 h, gently agitating every 10 min. We stored the lysed samples at -80 °C until further processing.

After thawing the pre-processed RNA samples on ice, we transferred the lysate to Lysing Matrix C tubes (MP Biomedicals, Santa Ana, CA, USA) that contain 1 mm silica beads. For a subset of samples (tRNA sample IDs: 1–12), we used Biosphere® SC Micro Tubes (Oakland, CA, USA) with 100 mg of 1 mm silica lysing beads for bead beating. We added 400 µL of 0.3 M NaOAc/HOAc, 10 mM EDTA, pH4.8 and an equal volume of acetate-saturated phenol/chloroform and vortexed briefly. We then placed the samples in a reciprocating bead beater for 2 min at maximum intensity, pausing after 1 min to avoid overheating the samples, and centrifuged at 13,000 x g for 15 min at 4 °C.

We transferred the aqueous phase to a fresh Lysing Matrix C tube and repeated the phenol/chloroform extraction steps. To precipitate the RNA, we added 1 volume of 100% isopropanol, vortexed each sample thoroughly, and incubated them overnight at -20 °C. After incubation we centrifuged the sample at 13,000 x g for 15 min at at 4 °C, discarded the supernatant, and washed each pellet with 1 mL of chilled 75% ethanol. Following centrifugation for 5 min at 13,000 x g at 4 °C and removal of the ethanol, we let the RNA pellet air dry for 10 min at room temperature to remove residual ethanol. We resuspended each pellet in 30 µL of an acid-elution buffer (10 mM NaOAc/1 mM EDTA, pH 4.8).

We cleaned the RNA using the Zymo Research Oligo Clean & Concentrate and eluted the RNA in RNAse-free water. Finally, we added an equivalent amount of acid-elution buffer to each extract to reach a final concentration of 5 mM NaOac/0.05 mM EDTA, pH 4.8. All samples yielded at least 200 ng of RNA except for tRNA sample 11.

### Sequencing of tRNA transcripts

We used approximately 200 ng of total RNA from each sample to build tRNA sequencing libraries following a previously published multiplex small RNA-seq (MSR-seq) protocol^25^. Following incubation at 37°C in a 33 mM sodium tetraborate buffer, pH 9.5 for 30 min to deacetylate RNAs, we added 5 µl of a polynucleotide kinase (PNK, New England Biolaboratories (NEB), USA) reaction stock (4 U/µL T4 PNK, 40 mM MgCl_2_, 200 mM Tris-HCl, pH 6.8) and incubated at 37 °C for 20 min to repair the 3’ end. We inactivated the PNK by incubating the samples at 65 °C for 10 min. We then added 30 µl of an RNA ligation reaction mix at a final concentration of 15% polyethylene glycol (PEG) 8000 (NEB, USA), 1× T4 RNA ligase I buffer (NEB, USA), 50 µM ATP, 5% DMSO, 1 mM hexaammine cobalt (III) chloride, and 1 U/µL T4 RNA ligase I (NEB, USA) and incubated the samples overnight at 16 °C. This mix also contained the barcoded RNA ligation linker/RT primer oligo at a 1.2:1 molar ratio to the input RNA to allow multiplexing of samples. After ligating the RNA to the barcoded and biotinylated hairpin oligonucleotide overnight, we added streptavidin-coated MyOne C1 dynabeads (ThermoFisher Scientific, USA) and incubated at room temperature on rotation for 15 min. This immobilized RNA molecules on the dynabeads in order to minimize sample loss and enabled rapid washes and buffer exchanges between reactions. To liberate the 3’ end of the reverse transcriptase (RT) primer by dephosphorylation, we added a phosphatase reaction mix (final concentration: 0.2 U/µL Calf Intestinal Phosphatase (CIP, NEB, USA), 10 mM MgCl_2_, 0.5 mM ZnCl_2_, 20 mM HEPES, pH 7.5) and incubated the samples for 30 min at 37 °C. To facilitate reverse transcription of the RNA molecules, we resuspended the samples in 25 µL of 1x SuperScript IV VILO (ThermoFisher Scientific, USA) mix and incubated at 55 °C for 10 min and then at 37 °C overnight. The following day, we added an RNase H reaction mix at a final concentration of 0.4 U/µL RNase H in 1× RNase H buffer (NEB, USA) for 15 min at 37 °C to digest the RNA molecules. To oxidize any non-extended reverse-transcriptase primer, we added a periodate solution at a final concentration of 50 mM sodium periodate in 15 mM sodium acetate, pH 5.0, and incubated the samples for 30 min at room temperature. We then ligated the barcoded cDNA ligation oligo to each cDNA molecule by incubating the samples overnight at room temperature in a 50 µL reaction with 2 U/µL T4 RNA ligase I (NEB), 25% PEG 8000, 7.5% DMSO, 50 µM ATP, 1 mM hexaammine cobalt (III) chloride, 2 mM barcoded cDNA ligation oligo and 1× T4 RNA ligase I buffer to enable polymerase chain reaction (PCR) amplification of the cDNA. Finally, we amplified the libraries via PCR with Illumina primers and performed multiplexed small RNA sequencing on an Illumina NovaSeq 6000 with 100bp paired-end sequencing. We used custom scripts to demultiplex the sequence data.

### Kāneʻohe Bay Time-series (KByT) collection and processing

To contextualize the diel data, we deeply sequenced four metagenomes collected at stations HP1 and STO1 on December 23, 2020 and May 24, 2021. These samples were part of the monthly sampling of the Kāneʻohe Bay Time-series and we processed the nucleic acids and other associated biogeochemical parameters following previously published protocols^18,19^. We followed the same short-read metagenomic library preparation and sequencing protocols as described above.

### Sequence data statistics and quality-control metrics

We exported read count and length statistics from the raw ‘omics data using ‘seqkit stats’ command in Seqkit v2.8.1^26^ with the ‘--all’ flag. To assess the quality of short-read metagenome and metatranscriptome libraries, we first examined the output of FastQC v0.12.1 (http://www.bioinformatics.babraham.ac.uk/projects/fastqc/). FastQC identified adapter sequences within the metatranscriptomic libraries, thus we used cutadapt v4.9^27^, with the following arguments to further trim adapter sequences and remove any subsequent reads with a length of less than 100 bp: ‘cutadapt -a AGATCGGAAGAGCACACGTCTGAACTCCAGTCA-A AGATCGGAAGAGCGTCGTGTAGGGAAAGAGTGT -n 5 -o trimmed-R1.fastq –p trimmed-R2.fastq input-R1.fastq input-R2.fastq -m 100’. We then used sortMeRNA v4.3.7^28^ with the ‘paired_out’, ‘fastx’, and ‘out2’ flags to assess the percentage of reads aligning to ribosomal RNA databases, specifically the SILVA v119^29^ silva-arc-16s-id95, silva-arc-23s-id98, silva-bac-16s-id90, silva-bac-23s-id98, silva-euk-18s-id95, and silva-euk-28s-id98 databases and Rfam 11.0^30^ rfam-5.8s-database-id98 and rfam-5s-database-id98. Next, we filtered the reads that did not match to rRNA databases using the illumina-utils library v1.4.1 called ‘iu-filter-quality-minochè^31^, which uses quality filtering parameters previously described^32^. We then evaluated the proportion of tRNA-sequencing transcripts that had features of tRNAs using ‘anvi-trnaseq’ from anvi’o v8.0^33^. Briefly, sequences were profiled from the 3’ end for base-paired stems of the proper length and a group of canonically conserved nucleotides, requiring the presence of a characteristic set of nucleotides that form and extend through the anticodon loop for a read to be identified as having a tRNA origin.

### Metagenomic co-assembly reconstruction and profiling

For both short- and long-read metagenomic data from HaDS, we co-assembled the data by grouping samples into those collected at coastal station HP1 and those collected at offshore station STO1 during the diel study. For the short-read metagenomic data, we used a Snakemake^34^ metagenomic workflow in anvi’o v8.0^33^ with the ‘anvi-run-workflow’ command and the ‘metagenomics’ flag. Briefly, the workflow quality controlled metagenomic short reads with illumina-utils^31^ and co-assembled metagenomic samples with idba_ud (https://github.com/loneknightpy/idba). Subsequently, we created anvi’o contig databases for each co-assembly with ‘anvi-gen-contigs-databasè, recruited metagenomic reads from all diel short-read metagenomes to the co-assemblies with bowtie2 v2.5.4^35^, and linked the read recruitment results back to the contigs database using ‘anvi-profilè. We further binned the co-assembled data using Metabat2 v2.12.1^36^, and imported these binned data into anvi’o as a collection with the command ‘anvi-import-collection’. To identify the taxonomic identity of the metagenomic bins, we ran ‘gtdbtk classify’ using the Genome Taxonomy Database Toolkit (GTDB-Tk) v2.4.0^37^. We used the ‘lineage_wf’in CheckM v1.2.3^38^ to assess the contamination and completion of the binned data and the ‘seqkit stats’ command in Seqkit v2.8.1^26^ with the ‘--all’ flag to examine total genome length, %GC, and N50 scores.

We co-assembled long-read metagenomes with Hifiasm v. 0.13-r308 ^39^ using default parameters. We split the largest contigs (>1 Mbp) from the co-assembly fasta file to individual fasta files to examine their quality and taxonomic identity using the ‘seqkit split’ command in Seqkit v2.8.1^26^. For each large contig, we built contig databases in anvi’o and assigned taxonomic identities and examined quality metrics as described above for the short-read metagenomes.

To evaluate the percent of reads mapped to each of the four coassemblies, we used samtools v1.21^40^ to extract information from mapping files. We mapped the long-read metagenomes back to the co-assemblies with minimap2 v2.28 (r1209)^41^ using the ‘--split-prefix’ and ‘-ax map-hifì flags. We mapped all quality-controlled short-read metagenomes and metatranscriptomes to the co-assemblies with bowtie2 v2.5.4^35^.

### Taxonomic profiling of short-read metagenomes, metatranscriptomes, and tRNA transcripts

Using quality-controlled short-read metatranscriptomes and metagenomes, we profiled the taxonomic composition of the reads using kaiju v1.10.1^42^ and the nr_euk database, which includes a subset of the NCBI BLAST nr database of proteins belonging to archaea, bacteria, and viruses, in addition to proteins from fungi and microbial eukaryotes. We summarized taxonomic profiles at the class level using the ‘kaiju2tablè command with the ‘-r class’ flag. For the tRNA-sequencing reads that were identified as tRNA transcripts, we dereplicated profiled tRNA sequences across samples using ‘anvi-merge-trnaseq’, yielding a set of tRNA seed sequences.

Seeds were taxonomically classified by ‘anvi-run-trna-taxonomy’ via alignment to tRNAs identified by tRNAscan-SE v2.0^43^ in the prokaryotic genomes of the GTDB database (release 95)^44^. Finally, we used ‘anvi-tabulate-trnaseq’ to output summary tables of tRNA abundances and taxonomy.

### Table and figure production

We plotted figures using R v 4.4.1^45^ and ggplot2 v 3.5.1^46^. We mapped sampling locations of HaDS and KByT using R with ‘geom_sf’ from ggplot2 based on geospatial data of the main Hawaiian Islands (USGS Digital Line Graphs). To draw contours, we accessed bathymetry data hosted by NOAA using marmap v 1.0.10^47^. Figures were edited in Inkscape v1.3.2 (https://inkscape.org/).

### Data Records

Metadata are available on the Simons Foundation Collaborative Marine Atlas Project (CMAP)^48^ and at the NSF Biological and Chemical Oceanography Data Management Office (BCO-DMO) under Project 947647^49^, in addition to **Table S1**. All sequencing products can be assessed through BioProject ID PRJNA1201851^50^ hosted by NCBI with SRA IDs available in **Table S1**. High-quality short-read metagenome assembled genomes (MAGs) are available under BioProject ID PRJNA1235278^51^ with accessions JBMHBP000000000-JBMHDZ000000000.

The remaining long-read and medium-quality short-read MAGs are available on FigShare^52^. Short-read and long-read metagenomic coassemblies are also available on FigShare^53^.

### Technical Validation

For short-read metagenomic libraries, we evaluated the DNA concentration using the Qubit dsDNA high-sensitivity (HS) assays (Thermo Fisher Scientific) and estimated the purity with a NanoDrop Microvolume Spectrophotometer (Thermo Fisher Scientific) before library preparation. We quantified the short-read metagenomic libraries using real-time PCR using KAPA library quantification reagents (Roche 07960204001) prior to sequencing. For DNA used in long-read metagenome sequencing, we quality-controlled DNA using HS Qubit assays (Thermo, Waltham, USA) and pulse-field capillary electrophoresis (Agilent FEMTOpulse). We evaluated the quality of RNA that served as input for metatranscriptome libraries with HS Qubit assays (Thermo, Waltham, USA) and a PicoChip (Agilent Bioanalyzer) for total RNA. Prior to transfer RNA sequencing library preparation we evaluated the quality and quantity of the RNA with the Qubit RNA HS assay (ThermoFisher Scientific, USA) and a Nanodrop Microvolume Spectrophotometer (ThermoFisher Scientific, USA).

Initial processing of HaDS raw sequences further confirmed the quality of the data. We filtered short-read metagenomes using illuminia-utils^31^, and retained 94.3±0.6% of reads (n=69; mean±sd; **Table S1**). We used cutadapt v4.9^27^ and sortMeRNA v4.3.7^28^ to remove reads matching to adapter sequences and ribosomal RNA (rRNA) databases, and then filtered low-quality reads from paired-end sequences^31^. The libraries retained 74.8±6.7% of the original paired reads and only a small fraction matched to rRNA databases (9.7±7.8; n=59; **Table S1**). These proportions demonstrate effective rRNA removal during library preparation and suggest that a large fraction of reads represent messenger RNA, underscoring their utility for gene expression analyses. We used ‘anvi-trna-seq’ within anvi’o v8.0^33^ to identify the proportion of reads that had characteristics of tRNA sequences. On average, tRNA transcripts accounted for 24.1±14.8% of the merged tRNA-sequencing reads, equating to an average of 1,871,536±2,931,481 tRNA transcripts per library (n= 66; **Table S1**). These proportions are in line with previous tRNA-sequencing experiments of the gut microbiome of mice (22.9±13.7%; n=6)^54^.

Taxonomic profiles of the quality-controlled short-read metagenomic, metatranscriptomic, and tRNA sequences showed communities consistent with the expected composition of coastal and offshore marine microbial communities from the oligotrophic tropical Pacific (**Fig. 2**; **Fig. S1**). Reads unclassified at the phylum-level or above were common among all three sequencing types (tRNA libraries: 33.14±7.26%, n= 66; metagenomes: 36.61±3.89%, n=65; metatranscriptomes: 66.02±8.50%, n=59). These proportions reflect known limitations in the sensitivity and specificity of short-read taxonomic profiling^55^, particularly in complex and under-characterized microbial communities. *Alphaproteobacteria* and *Cyanophyceae* dominated short-reads classified at the phylum-level or below (**Fig. 2, Table S1**), as expected for surface ocean marine communities^56^. All three sequencing types captured differences in community composition between the two stations, with *Mamiellophyceae* and *Flavobacteriia* elevated in coastal samples (**Fig. 2, Table S1**), as previously described^18,19^. Shifts in community composition between day and night samples were evident in coastal metagenomes, metatranscriptomes, and tRNA transcripts, and present although more subtle in the offshore sequences (**Fig. 2, Fig. S1**). The relative abundances of the dominant taxa broadly aligns across sequencing types, suggesting that future integrated data analysis has the potential to reveal drivers of microbial physiologies across time and space.

**Figure 2.**
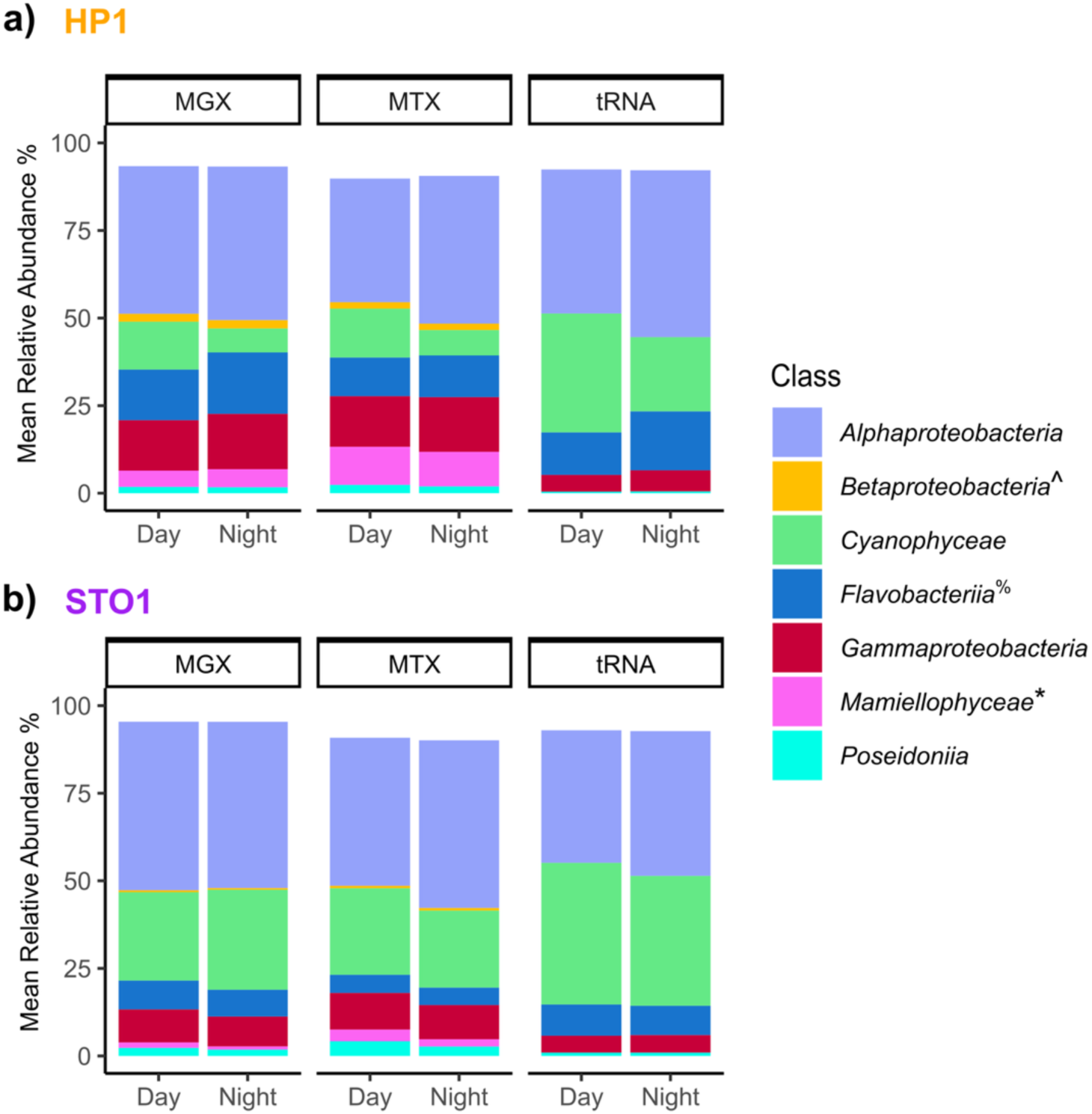
Taxonomic profiling across sample types and stations. We profiled the taxonomy of quality-controlled, paired-end metagenomic (MGX) and metatranscriptomic (MTX) reads and tRNA transcripts for coastal HP1 (**a**) and offshore STO1 (**b**). We averaged the relative abundances for classes that represent >1% mean abundance of the total community across samples collected during sunlit hours (day) and those collected after sunset (night). The bar chart shows the fraction of reads with assignments resolving to at least phylum-level taxonomy. The identification of tRNA sequences was based on the GTDB database, and the identification of metatranscriptomic and metagenomic reads was based on the nr_euk database that utilizes taxonomy from NCBI. *Mamiellophyceae: GTDB does not identify eukaryotic organisms. ^%^Flavobacteriia: Flavobacteriia is an NCBI class and an order in GTDB (c Bacteroidia; o Flavobacteriales). For tRNA sequences, the GTDB Bacteroidia class was dominated by reads matching to the order Flavobacteriales. ^Betaproteobacteria: Betaproteobacteria is not its own class within GTDB, and is part of the class Gammaproteobacteria.

A high proportion of short-reads mapped to the respective station-specific short-read and long-read co-assemblies, with 67% to 82% of short reads mapped to assemblies on average (**Table S1**). Similarly, a high proportion of long-read libraries mapped to the respective station-specific short-read and long-read co-assemblies, where 61% to 75% of the total length of the long-read library mapped to the station-specific assembly on average (**Table S1**). The station-specific short-read and long-read co-assemblies recruited a smaller yet still substantial percentage of the metatranscriptomic reads (33-60% of reads mapped on average; **Table S1**). To ensure the quality of the metagenome assembly genomes (MAGs), we examined N50 metrics and completeness and contamination scores (**Table S1**), and only reported those of high and medium quality, >90% completion and <5% contamination or >70% completion and <10% contamination, respectively. Among the high-quality long-read MAGs is a circularized contig belonging to *Pelagibacterales* (SAR11), a prevalent marine microbial clade with genomes that are notoriously difficult to reconstruct from short-read metagenomes^57^. While recent work highlighted pervasive instances of misreporting of circular contigs from long-read assemblers^58^, our manual inspection of this genome suggests that it does not suffer from the kinds of issues (such as extensive read-clippings and excessive repeats) previously reported^58^.

Finally, we cross-referenced and contextualized the biogeochemical observations from the HaDS dataset with long-term environmental data from nearby buoys and weather stations (**Fig. 1**; **Fig. S2**), as well as published values from the Kāneʻohe Bay Time-series^18,19^.

### Usage Notes

#### Biogeochemical context of HaDS

The environmental variation sampled across space and time offers valuable opportunities to explore how microorganisms respond to biotic and abiotic factors. Stations HP1 and STO1 differ in their physical characteristics and biogeochemistry^18,19^. During HaDS, station HP1 was characterized by warmer waters, lower salinity, and elevated phytoplankton biomass and inorganic nutrient concentrations relative to STO1, as expected during the summer months (**Fig. 3a; Table S1**). These environmental differences align with shifts in the microbial community, such as higher abundances of photosynthetic picoeukaryotes, *Syncechococcus*, and heterotrophic bacteria at station HP1 compared to the offshore environment. In contrast, offshore station STO1 is enriched in *Prochlorococcus* cells relative to coastal Kāneʻohe Bay (**Fig. 3a**).

**Figure 3.**
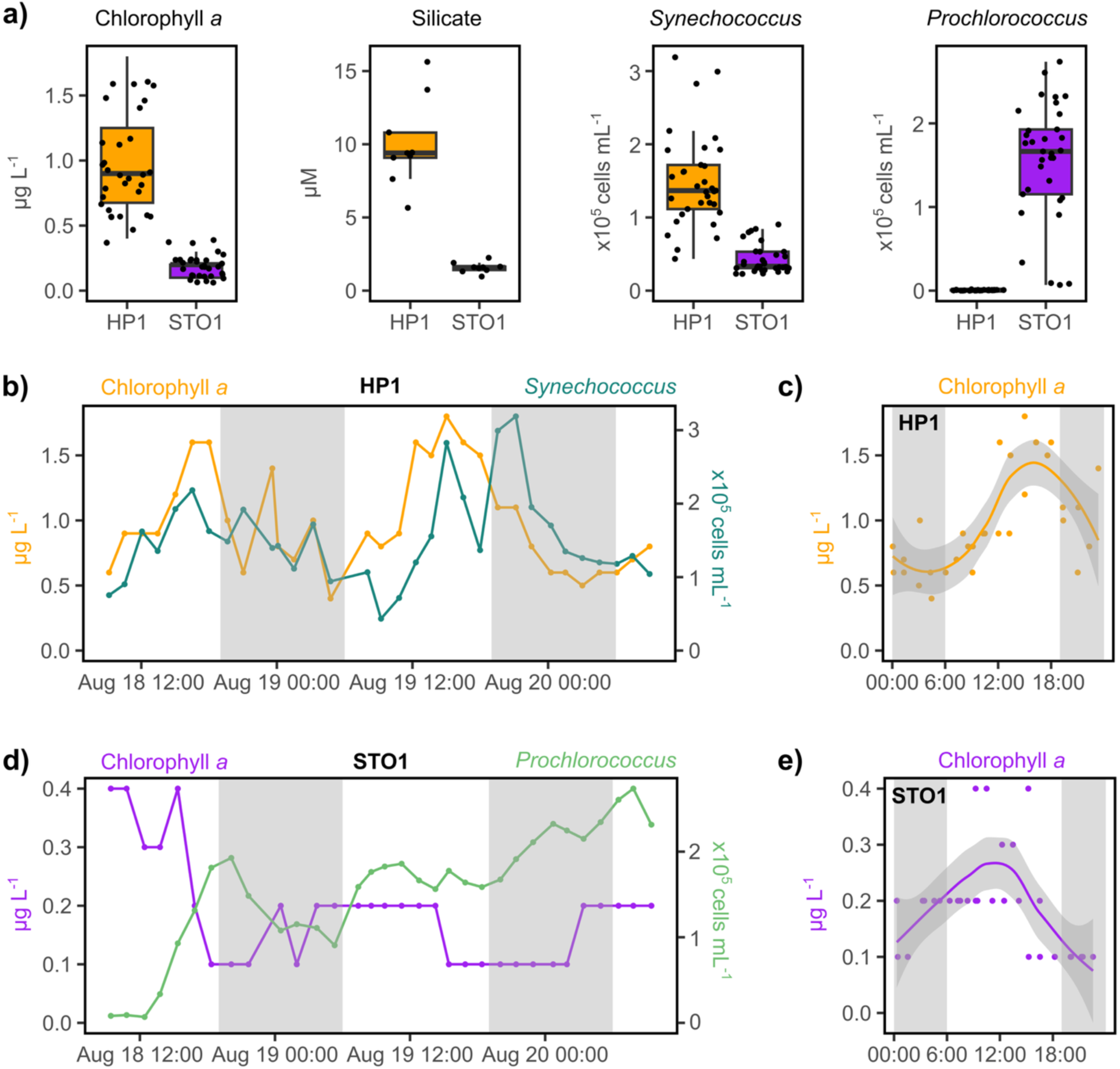
Variation in biogeochemistry and phytoplankton communities across space and time during HaDS. **a**) Concentrations of chlorophyll *a* and silicate and *Synechococcus* cellular abundances are higher at coastal station HP1 compared to offshore station STO1, while *Prochlorococcus* cells are more abundant at offshore station STO1. **b**) At station HP1, chlorophyll *a* concentrations and *Synechococcus* cellular abundances follow diel oscillations, with early morning (∼03:00 HST) minimums and late afternoon (∼15:00-21:00 HST) maximums. Chlorophyll *a* concentrations at station HP1 rise during daylight hours and decline after sunset. Samples are plotted by time of day over a 24 h cycle. **d**) At station STO1, *Prochlorococcus* cellular abundances show a clear trend that emerges on the order of days while chlorophyll *a* concentrations decline briefly at the beginning of the sampling period. **e**) Chlorophyll *a* concentrations at station STO1 reached a minimum in the late evening. Like panel c, this plot shows samples by time of day over a 24-hour cycle. The shaded areas in panels c and e depict the standard error around a local polynomial regression fit line for chlorophyll *a* concentrations per hour of the day. Gray bars in panels b-e indicate nighttime.

Over the 48 h sampling period, HP1 experienced temporal patterns that likely reflect a diel signal (**Fig. 3b, c**). Although station STO1 also exhibited subtle diel patterns, the dominant temporal trend emerged over several days where, throughout the 48-hour diel sampling, chlorophyll *a* concentrations declined while *Prochlorococcus* cell numbers increased (**Fig. 3d, e**). The dominant temporal pattern at STO1 likely reflects broader climatic conditions that developed over the days leading up to the diel sampling (**Fig. S2**). Wind and wave-driven currents typically transport offshore waters into the bay across the centrally-located barrier reef. Variations in wind speeds and wave energy can strongly impact the circulation patterns that influence nearshore to offshore gradients both within and adjacent to Kāneʻohe Bay^59^.

Additionally, nutrient-enriched stream discharge following heavy rain events can trigger rapid phytoplankton responses in the bay^19,60–62^. The relatively subdued onshore wind and wave height prior to the diel sampling, in combination with a brief but intense rainfall event during an outgoing tide shortly before sampling (**Fig. S2**), may have facilitated the export of nearshore waters from coastal Kāneʻohe Bay to offshore station STO1 and temporarily altered the biogeochemical conditions at STO1.

#### Label convention

Labels follow a standardized naming convention to enable cross-referencing with relevant environmental data and between ‘omic data types. For example, we have sample file HADS_20210818_H1030_MGX_STO1_003_R1_001.fastq.gz

where each of the following fields are separated by an underscore:

- Project name: HaDS stands for Hawaiʻi Diel Sampling.
- Sampling date: Four digit year, two digit month, and two digit day, where day restarts at midnight.
- Sampling hour: Starts with letter H, military time for hours and minutes with no other characters in between. Times given are in Hawaiʻi Standard Time.
- Sample type: MGX (short-read metagenomes), MTX (metatranscriptomes), TRN (tRNA-seq), or HMW (long-read metagenomes from high-molecular weight DNA).
- Sampling station: Can be STO1 or xHP1. Note that although we use xHP1 in the sample name to represent the station HP1 the formatting is solely to create equal-length names to facilitate computation, and reference to the station should continue to explicitly use HP1, not xHP1.
- Sample ID. This number is unique within each sample type.

The following three fields may be omitted from sample names that appear in analyses, but must always be a part of the file names of raw data:

- Read. R1 or R2. Only relevant for data files.
- Sequencing run (in case there is more sequencing data for the same sample and data type). If multiple sequencing runs were concatenated to one file, then this field in the concatenated file will be 000.

#### Metatranscriptomes

We recommend additional removal of adapter sequences and the removal of short reads (<100 bp) prior to use of the metatranscriptomic libraries, as outlined in the methods section.

## Code Availability

The URL https://merenlab.org/data/HaDS gives access the bioinformatics workflow we implemented to perform quality control, metagenomic co-assembly, read mapping, and recovery of metagenome-assembled genomes, as well as the reproducible anvi’o data artifacts^52,53^.

## Acknowledgements

We thank Jason Jones and Andrew Brown for their assistance with field operations. The data collection and curation was supported by the W. M. Keck Foundation, the Simons Foundation (grant #687269), and the Center for Chemical Currencies of a Microbial Planet (C-CoMP) (NSF grant OCE-2019589) to AME and the Simons Foundation to SJT (grant # 989028). This is C-CoMP publication XXX, SOEST contribution XXX, and HIMB contribution XXX. Data provided by PacIOOS (www.pacioos.org), which is a part of the U.S. Integrated Ocean Observing System (IOOS), was funded in part by National Oceanic and Atmospheric Administration (NOAA) Awards #NA16NOS0120024 and #NA21NOS0120091.

## Author Contributions

Conceptualization: SJT, JF, KCF, TP, MSR, AME

Methodology: SJT, JF, KCF, MS, SEM, FT, KL, HGM, BF, BH, TP, MSR, AME

Investigation: SJT, JF, KCF, EK, EBF, OR, CESS, APG, HM, MRS, MQ, CR, LLT, MS, SEM, FT, KL, HGM, BF, BH, TP, MSR, AME

Visualization: SJT, AME Supervision: MSR, AME

Writing—original draft: SJT, JF, KCF

Writing—review & editing: SJT, JF, KCF, MSR, AME

