## Supplementary Information for "A high-resolution diel survey of surface ocean metagenomes, metatranscriptomes, and transfer RNA transcripts"

**This PDF file includes:**

Figs. S1 and S2

**Other Supplementary Information for this manuscript include the following:**

Table S1 (Excel Spreadsheet)

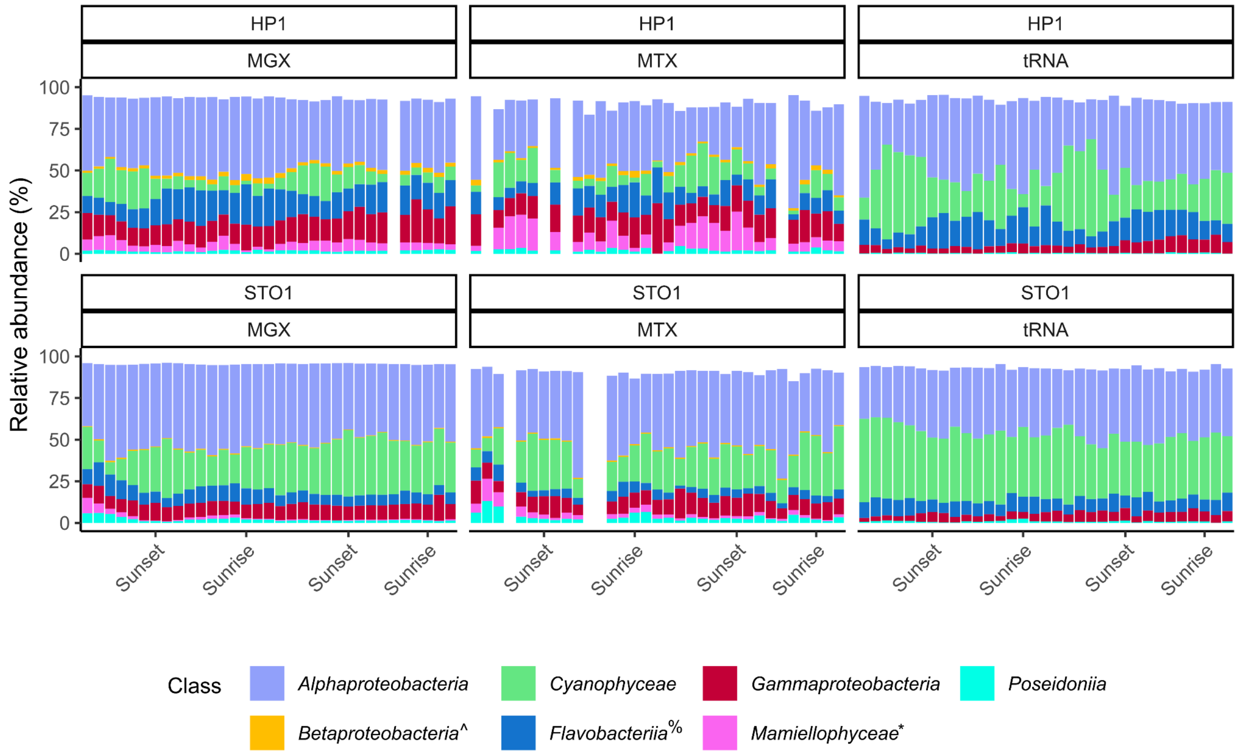


**Fig. S1. Taxonomic profiling across samples.** A comparison of the taxonomic profiles for quality-controlled, paired-end metagenomic (MGX) and metatranscriptomic (MTX) reads and tRNA transcripts across all samples. Samples are chronologically ordered based on the time of collection, with sampling events coinciding with sunrise and sunset noted. We show taxonomic classes that represent >1% mean abundance of the total community in at least one sample type. Only the fraction of reads that could be assigned to phylum-level taxonomy are included. The tRNA sequences were identified with the GTDB database, while the metatranscriptomic and metagenomic reads were identified with the nr_euk database that utilizes taxonomy from NCBI. *Mamiellophyceae: GTDB does not identify eukaryotic organisms. %Flavobacteriia: Flavobacteriia is an NCBI class and an order in GTDB (c__Bacteroidia; o__Flavobacteriales). For tRNA sequences, the GTDB Bacteroidia class was dominated by reads matching to the order Flavobacteriales. ^Betaproteobacteria: Betaproteobacteria is no longer a class within GTDB, and is now part of the class Gammaproteobacteria. Empty bars signify that sequence data are missing for that data point.

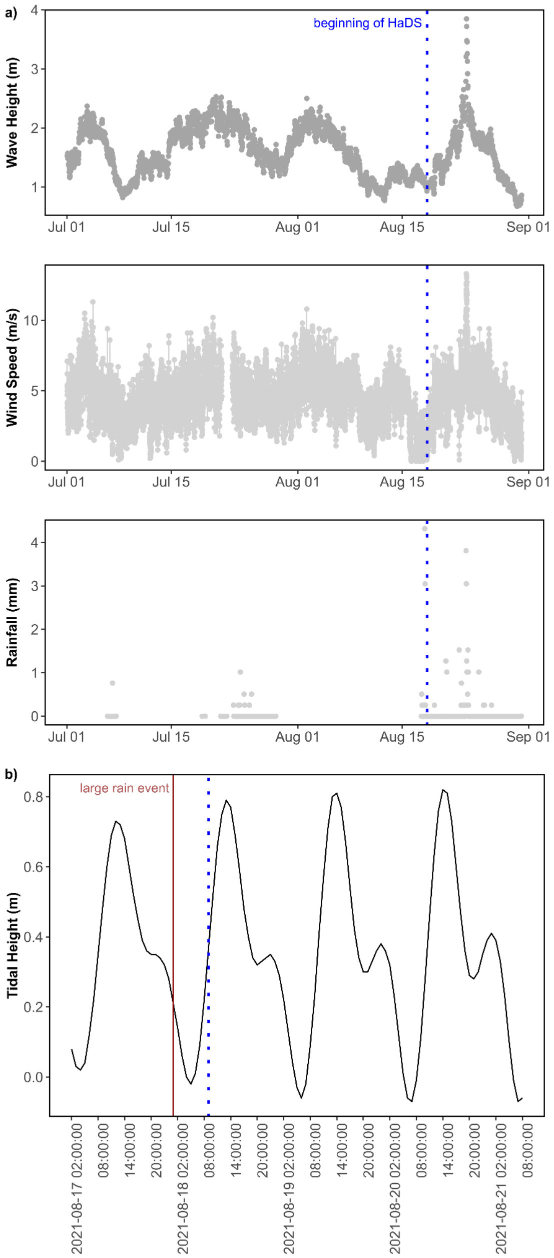


**Fig. S2. Weather and oceanographic characteristics leading up to HaDS.** Wave height, wind speeds, and rainfall were plotted from July 1, 2021, to September 1, 2021, to show the month before HaDS (dotted blue line) (**a**). Wave height and wind speed dropped immediately before the diel sampling. Heavy rain occurred at 1:00 AM Hawaiʻi Standard Time (HST) on August 19, 2021, the early morning immediately before the diel sampling started at 9:00 AM, with a second lighter rain event also occurring during the diel sampling (August 20, 2021 at 21:00 PM HST). The heavy rainfall event occurred during an outgoing tide (**b**). Tidal height is plotted from August 17, 2021, to August 21, 2021, to show the day before the sampling event. Wave height was collected by offshore NOAA buoy (station 51207h), weather data (wind speed and rainfall) were collected by the Moku o Loʻe Weather Station through NOAA's Pacific Islands Ocean Observing System, and tidal height was collected by NOAA/NOS/CO-OPS Station ID 1612480.
